# In-silico analysis of Salmonella typhimurium and E.coli Methionyl tRNA synthetase at primary, secondary, tertiary level with protein disorder and functional association of differences

**DOI:** 10.1101/048009

**Authors:** Prabhakar B. Ghorpade, Pooja Kadu, Amit Pandey, Yogesh Banger, Bhaskar Sharma

**Affiliations:** College of Veterinary and Animal Sciences, Parbhani

**Keywords:** Methionyl-tRNAsynthetase, Protein Disorder, Primary Structure, Secondary Structure, Tertiary Structure, HomologyModelling, Superimposition, Docking

## Abstract

The identification changes in amino acid for same protein in closely related species are necessary in order to identify its effect at various structural and functional levels. Salmonella typhimurium and E.coli Methionyl tRNA synthetase taken in current study as these bacteria are closely related to each other and have fewer differences in amino acid sequences for MetG. This study helps to identify various structural and functional differences at primary, secondary and tertiary levels, with functional differences by Docking study with Methionine. Study involves analysis of differences based on observation of differences in modeled 3D protein for sequences available at NCBI and its comparison with Known 3D structure. As sequences difference are in functional protein from non-mutant species, the differences are analysed in context of Primary, secondary, tertiary structure differences, Disorder differences, and docking differences.

## Introduction

Aminoacyl-Trna synthetase catalyzes the binding of an amino acid to 3- end of tRNA by using the energy generated by dephosphorylation of ATP. There are 20 aminoacyl tRNA synthetases which are differentiated into two classes, Class I and Class II. Each class contains 10 aminoacyl tRNA synthetase. Methionyl-tRNA synthetase (EC 6.1.1.10) is a class I aminoacyletRNAsynthetase and catalyze the ligation of methionine to tRNA Met; the whole reaction can be represented as:

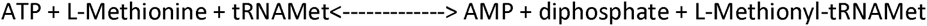

Methionyl-tRNA synthetase (MetG) recognizes both initiator tRNA and methionine tRNA. Like all class I aminoacyl tRNA synthetase, MetG has acatalytic domain and an α helix bundle/anticodon binding domain. Catalytic domain consist of Rossman Fold, Stem Contact (SC) Fold and CP (Connective Polypeptide)/Zinc binding domain (Deniziak and Barciszewski, 2001). Rossman fold has two Class I signature motifs “HIGH” and “KMSKS”. SC fold helps in contact of MetG with tRNAMet (Sugiura, et al., 2000). Zinc finger helps methionine activation and correct positioning of 3’ end of tRNA during translation(Fourmy, et al., 1995).

Aminoacyl tRNA synthetases perform similar function across different species. Their amino acid sequences are highly conserved across species. Crystal structure of many aminoacyl tRNA synthetases are available making it easier to predict 3 D structure of corresponding aminoacyl tRNA synthetase from other closely related species. We are currently working on MetG (Methionyl-tRNA synthetase) of Salmonella typhimurium (STM) an important serovar of species Salmonella enetrica sub species enterica, which has zoonotic importance. STM cause disease in humans and animals; horizontal transfer through egg to human has also been reported. Our current works on MetG of STM involves generating mutants for structural functional studies. As a prelude to that in this study we have compared 3D predicted structure of MetG of Salmonella and E.coli taking crystal structure of metG of E.coli as template to check whether naturally evolved and closely related proteins reflect the difference in amino acid sequences in their 3D structure. We have superposed both the predicted structure and also the predicted structure with crystal structure and report the results.

Identification of difference between protein sequences available at NCBI is major challenge. To know difference of the partial and full length proteins sequences of Methionyl tRNA synthetase of *E.coli* and *Salmonella typhimurium* were analysed. As E.coli and Salmonella typhimurium are both proteobacteria both are expected to have similar Methionyl tRNA synthetase structures. Current study compared both full length protein structures with truncated. The protein used in study has been mentioned in Table No. 1. The amino acid differences between MetG and MetRS were noted in Table No. 2. The amino level differences mentioned in Table no. 2 hardly gives any interpretation of secondary structure difference and 3D structural difference. To solve mystery of amino acid compositional differences in protein interest as between same species (MetRS, MetRS551, 1QQT_A) or (MetG, MetG551) or between the interspecies E.coli and Salmonella typhimurium, we have undertaken this study. The differences are analysed at 1D, 2D, 3D and functional level using various available tools.

**Table 1.** Name of Protein structures:-

| Abbreviation |  | Full Name | Gene identifier |
| --- | --- | --- | --- |
| MetG | (Modeller determined structure) | Salmonella typhimuriumMethionyl-tRNA synthetase - 667 amino acids | gi\|16765484\|ref\|NP_461099.1\| |
| MetRS | (Modeller determined structure) | Escherichia Coli Methionyl-tRNA synthetase - 667 amino acids | gi\|485773692\|ref\|WP_001397492.1\| |
| MetG551 | (Swiss model determined structure) | Salmonella typhimuriumMethionyl-tRNA synthetase 2-552 amino acids | gi\|16765484\|ref\|NP_461099.1\| |
| MetRS551 | (Swiss model determined structure) | Escherichia Coli Methionyl-tRNA synthetase 2-552 amino acids | gi\|485773692\|ref\|WP_001397492.1\| |
| 1QQT_A | ( X-Ray Diffraction structure) | Chain A, Methionyl-TrnaSynthetase From Escherichia Coli 551 amino acids | gi\|6730520\| |

**Table 2a.**
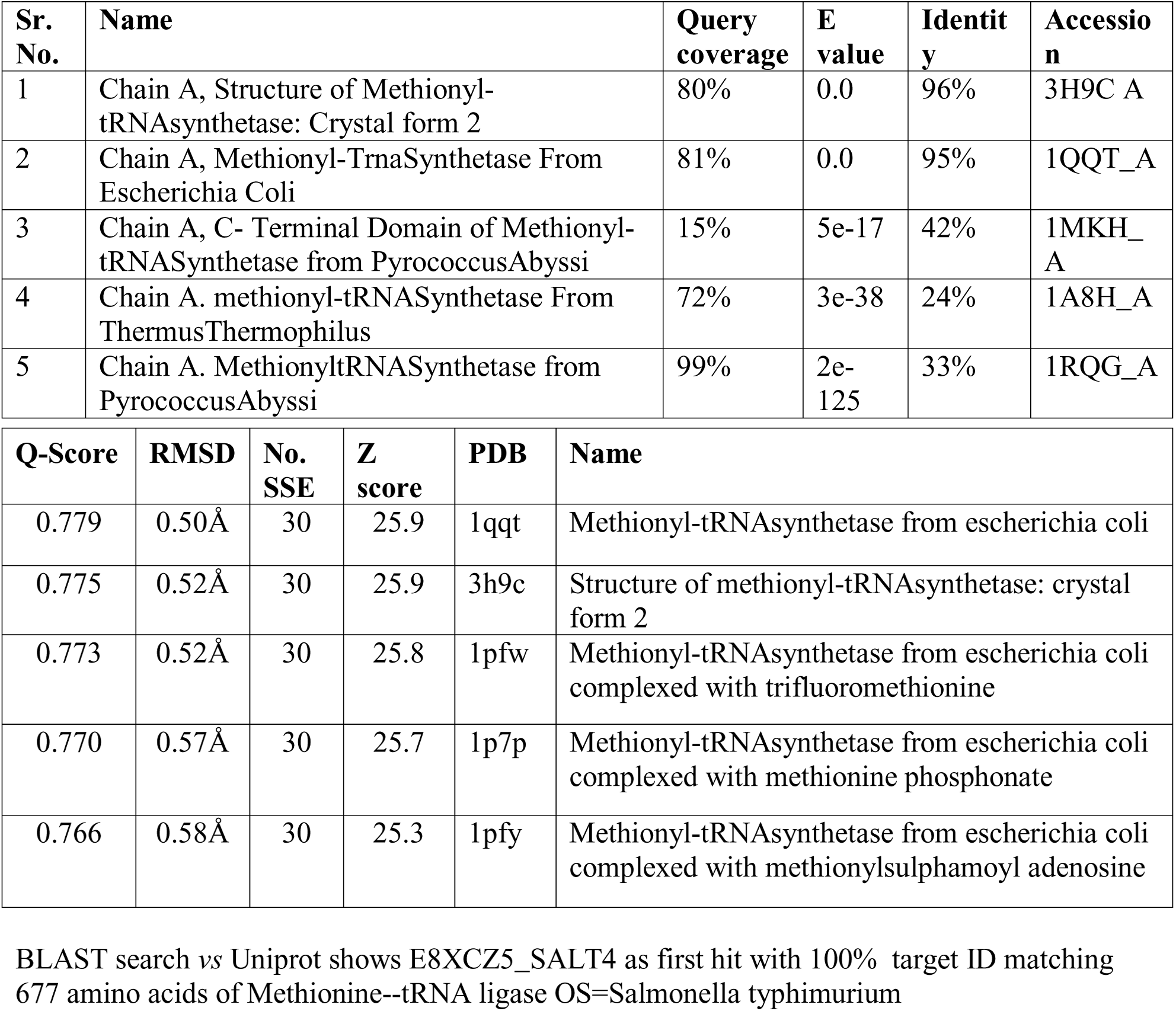
Template proteins selected for model building. b. Matching Fold for current MetGmodel from PROFUNC

The similar 3D structures are evolutionary more conserved than similar sequences. Taking this view three dimensional structures comparison could be more significant in predicting similarity in structures. Current investigation involves homology/ comparative modelling of salmonella typhimurium methionine trna-synthetase from available protein 3Dstructures in rcsbpdb (protein data bank). Then method modeller use for modeling is based on spatial restraint matching (Eswar, et al., 2006). In this amino acid on string of template was known how it folds, this folding is correlated with spatial restraints such as Cα-Cα distances, main chain and dihedral angles, hydrogen bonds. This correlation is used for predicting query sequence structure. The program uses known 3D structure of pdb as template and structure to be determined as query sequence. Apart from comparative modeling modeller can perform de-novo loop modeling, and optimising model based on objective functions.

## Material methods

### Sequences and sequence alignment

Salmonella (Ac.no. gi|16765484|ref|NP_461099.1|) and E.coli (AccNo. gi|485773692|ref|WP_001397492.1|) MetG amino acid sequences were down loaded from NCBI data base and were aligned using Clustal X to calculate the sequence similarity. Protein blast with query MetG sequence of Salmonella typhimurium against PDB database was done to find crystal structure similar to MetG. Selection of template for building the 3D structure was done on the basis of identity score and the structure without any bound ligand.

### Protein Compositional Differences

The compositional differences between protein was analyzed by protoparam.

### Phylogenetic Tree

The ClustalW aligned sequences were subjected to MEGA evolutionary analysis by Maximum likelihood method.

### Secondary Structure difference

The PDBsum was used for predicting secondary structure and Ramchandran plot parameter checking. (Laskowski, et al., 1993; Laskowski, et al., 2005).

### Protein Disorder Prediction

Protein exist in Stable ordered from, molten globule form and random coil form. The intrinsic disordered proteins lack fixed three dimensional shape. The binding of other proteins, DNA, RNA may facilitate the conversion of IDP to 3D structure imparting functional importance in molecular recognization, cell regulation and cytoskeleton structure. Current study involves disorder prediction for our genes of interest using Dismeta server(Huang, et al., 2014). The program or server for Disorder prediction of protein inculdes DISEMBL, DISOPRED2, DISpro, FoldIndex, Globeplot2, IUPred, RONN, VSL2. The all above mentioned programmes are combined in Dismeta and the individual plus combined result for them is presented. In Dismeta server addition to disorder prediction the identification of signal peoptide by signalIP; helix sheet by PROFsec, PSIPred and secondary structure consensus; Transmembrame segment prediction by TMHMM, low complexity prediction by SEG and protein binding site (disorder) by ANCHOR is done(Iakoucheva and Dunker, 2003; Linding, et al., 2003).

### Model Building

The template used in modeler for building MetG and MetRS models are mentioned in Table. No. 2. Model for MetG551 and MetRS were built using Swiss Model. The 1QQT_A model was taken as experimental model for comparison as Salmonella typhimurium MetG PROFUNC gives 1QQT_A as first hit. Detailed protocol for model building is as follows,

### Model Generation by Modeller

#### 1. Searching Structure related to toMetG

The Methionine tRNAsynthetase (MetG) protein of Salmonella typhimurium (NCBI Accession number:-NP_461099.1) was selected to build protein model of our interest.Modeller 9.12 (Eswar, et al., 2006) with Easy Modeller 4.0 as GUI(Graphical user Interface) was used for model building from template sequences. Query sequence MetG of Salmonella typhimurium is added, the five templates we selected based on Methionyltrnasynthetase without ligand from protein protein BLAST criteria were used for model building.

#### 2. Selecting template

The Protein-BLAST was performed against our query to search related sequences in NCBI against protein Data bank to search sequences showing homology to query sequence(Altschul, et al., 1997). BLOSUM 62 scoring matrix with default parameters were used in searching related sequences. The BLAST results show matching protein sequences for which 3D structure is available in PDB. Result in BLAST is based on scoring, Identity; we selected maximum identity as criteria to select template for modeling our query sequence structure. The Sole protein which is not complexedwith any chemical ligand is selected. Based on above mentioned criteria we selected five models in NCBI for structure prediction (Table. 1).

#### 3. Aligning MetG with template

The templates were compared among themselves based on Weighted pair-group average clustering based on a distance matrix then alignment of query with templates was done. Here query sequence alignment with available templates helps query sequence to take information such as spatial restraints from templates to build the model.

#### 4. Model building

Here number of Models to be built is specified, In our case we are taking 5 models to build. Model built is selected based on lowest DOPE score. For Viewing model Rasmol 2.7.5.2 was used. The Pymol(DeLano, 2002; DeLano, 2002) could be used for model visualization.

#### 5. Model Evaluation

Models produced was energy minimized in modeler by optimize option. For Model evaluation PROCHECK (stereochemical properties& Tertiary Structure analysis) (Laskowski, et al., 1993; Laskowski, et al., 2005), Verify 3D (Bowie, et al., 1991), Errata and Eval123D (For structural evaluations using EVAL23D, VERIFY3D, EVDTREE and ERRAT) were used. The Protein Analysis ToolKit (PAT) gives one spot model evaluation using multiple methods(Gracy and Chiche, 2005).

### Modelling by Swiss Model-

In this sequence of amino acid was submitted to swiss model workspace and automatic mode model was generated for MetG551 and MetRS551(Arnold, et al., 2006; Schwede, et al., 2003).

### Experimental Model

Crystal structure 1QQT_Awas searched in PDB database and down loaded from it(Berman, et al., 2002).

### Visualisation of 3D structure

Rasmol and PyMol were used to visualize the model.

### Superposition of Protein Structures

The 3D structures are more evolutionary conserved as compared to sequence of protein. The SuperPose web server we used in current study uses combination of various parameters as sequence alignment, difference distance matrix comparison, and quaternion eigenvalue superposition. Superpose is used to find structural differences in protein. The combinatorial extension was also used for superimposition (Maiti, et al., 2004).

### Swiss Docking

The Swiss docking was performed on each five structures with L-Methionine (ZINC01532529) as ligand.

## Results

Overall steps followed in current research are mentioned in Fig. 1

**Fig1.**
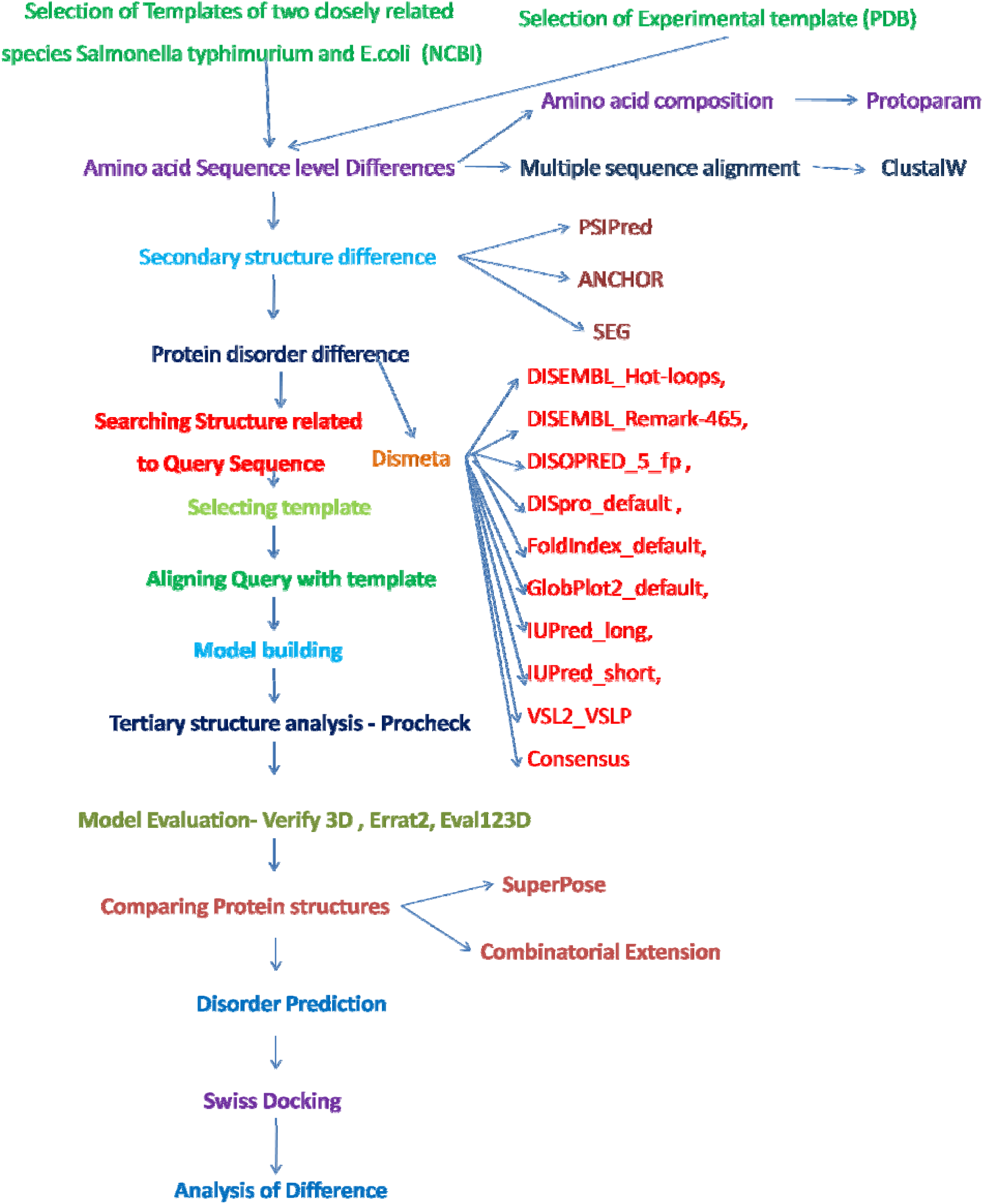
Flow Diagram of Protein Modeling and Structure Comparison

The current study uses full length and partial length Methionyl-tRNA-synthetases from Salmonella typhimurium (MetG) (Fig2.), E.Coli (MetRS), and experimentally determined partial structure 1QQT_A (Table 1).

**Fig2a.**
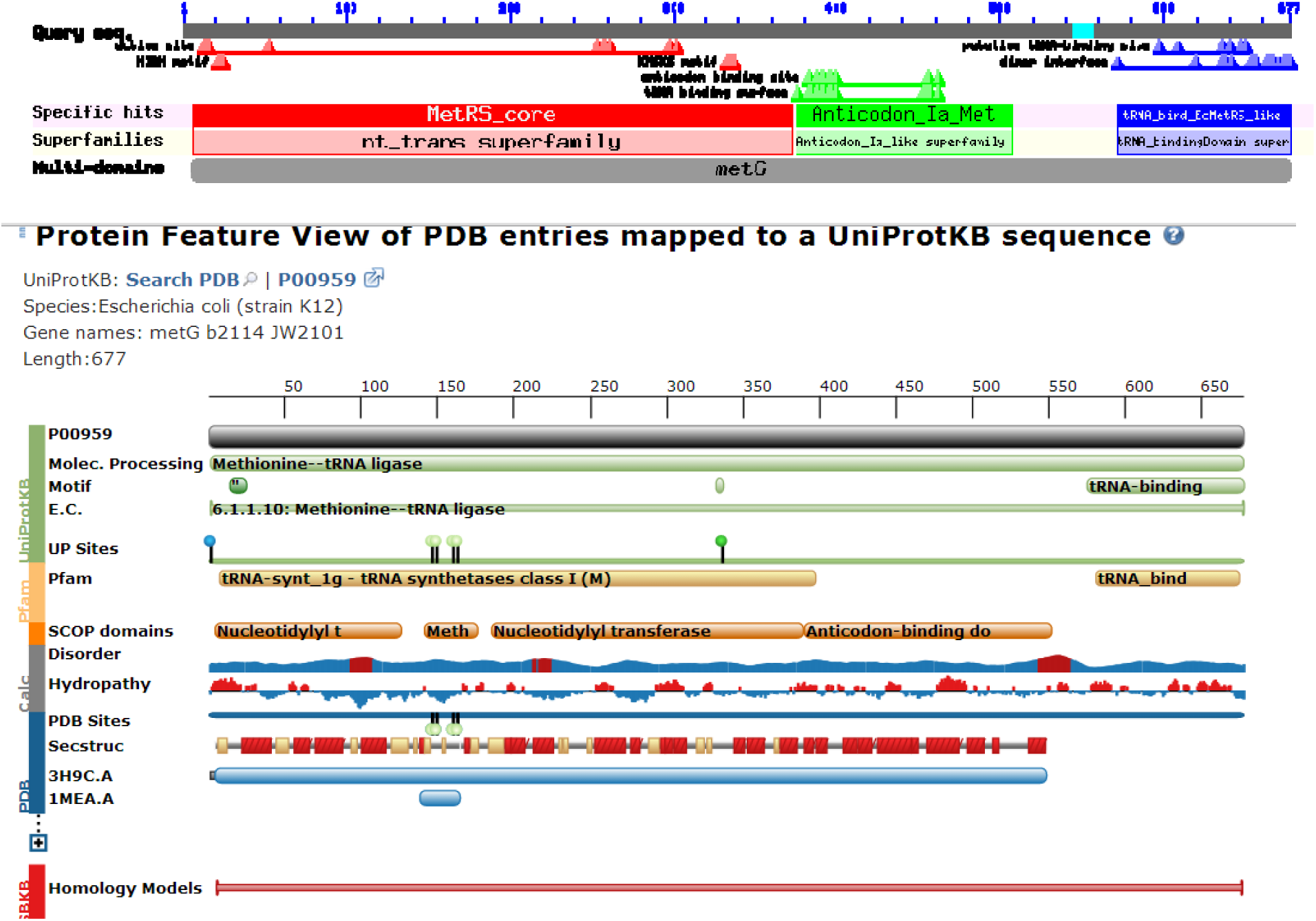
MetG Protein superfamilies from NCBI b. NCBI Protein Feature View of Methionyl-tRNAsynthetase

Amino acid Sequence level Differences-The protein sequences of interest were compared at based on compositional basis and multiple sequence alignment. Sequences MetG and MetRS shows differences in composition of amino acids as Ala, Asn, Asp, Glu, Gly, His, Ile, Leu, Lys, Phe, Pro, Ser and Thr (Table3, Table 4). Multiple alignment showing difference in composition of proteins are shown in figure 3. The position specific sequence feature forMethionyltRNAsynthetase are shown in Table. No.5. The visual comparison of aligned sequences will help to identify sites mutation which would not be having difference on functional aspect of protein as both sequences were retrieved from public database (NCBI) with non-mutant sequences of E.coli and Salmonella typhimurium species. The mutational differences in amino acid sequences in intraspecies (E.coli) and interspecies (E.coli and Salmonella typhimurium) does not show specific differences in conserved regions or protein as depicted by Conserved domain search from NCBI(Bauer, et al., 2011; Marchler-Bauer, et al., 2009; Marchler-Bauer, et al., 2011). The position of group specific features can be found at Table.5 and Fig.2. Further as both sequences are natural bacterial sequences support claim that no significant difference in conserved domain are found possibly because of their essentiality to function of protein.

**Fig3.**
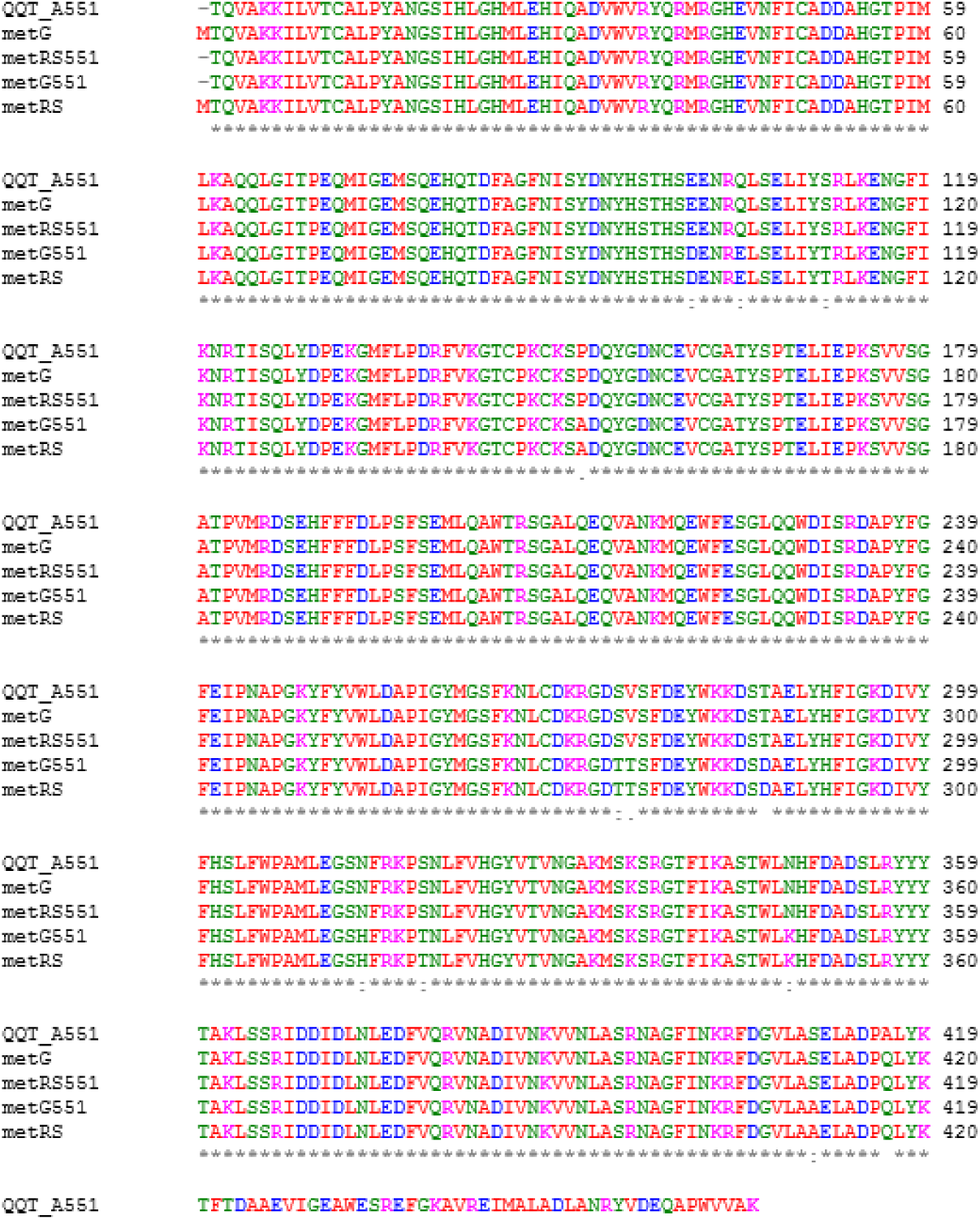
ClustalW Multiple Sequence alignment

**Table.3.** Amino acid composition of MetG Protein *Salmonella typhimurium* is as follows

| Name of amino acid | MetG Total a.a. number | MetGa.a. Percent | MetRS Total a.a. number | MetRSa.a. Percent |
| --- | --- | --- | --- | --- |
| **Ala (A)** | **60** | **8.9%** | **59** | **8.7%** |
| Arg (R) | 32 | 4.7% | 32 | 4.7% |
| **Asn (N)** | **28** | **4.1%** | **29** | **4.3%** |
| **Asp (D)** | **48** | **7.1%** | **45** | **6.6%** |
| Cys (C) | 8 | 1.2 | 8 | 1.2% |
| Gln (Q) | 28 | 4.1% | 28 | 4.1% |
| **Glu (E)** | **43** | **1.** | **44** | **6.5%** |
| **Gly (G)** | **45** | **6.6%** | **46** | **6.8%** |
| **His (H)** | **15** | **2.2%** | **16** | **2.4%** |
| **Ile (I)** | **37** | **5.5%** | **36** | **5.3%** |
| **Leu (L)** | **56** | **8.3%** | **57** | **8.4%** |
| **Lys (K)** | **41** | **6.1%** | **40** | **5.9%** |
| Met (M) | 21 | 3.1% | 21 | 3.1% |
| **Phe (F)** | **38** | **5.6%** | **37** | **5.5%** |
| **Pro (P)** | **30** | **4.4%** | **33** | **4.9%** |
| **Ser (S)** | **40** | **5.9%** | **41** | **6.1%** |
| **Thr (T)** | **30** | **4.4%** | **28** | **4.1%** |
| Trp (W) | 11 | 1.6% | 11 | 1.6% |
| Tyr (Y) | 24 | 3.5% | 24 | 3.5% |
| Val (V) | 42 | 6.2% | 42 | 6.2% |

**Table 4.**
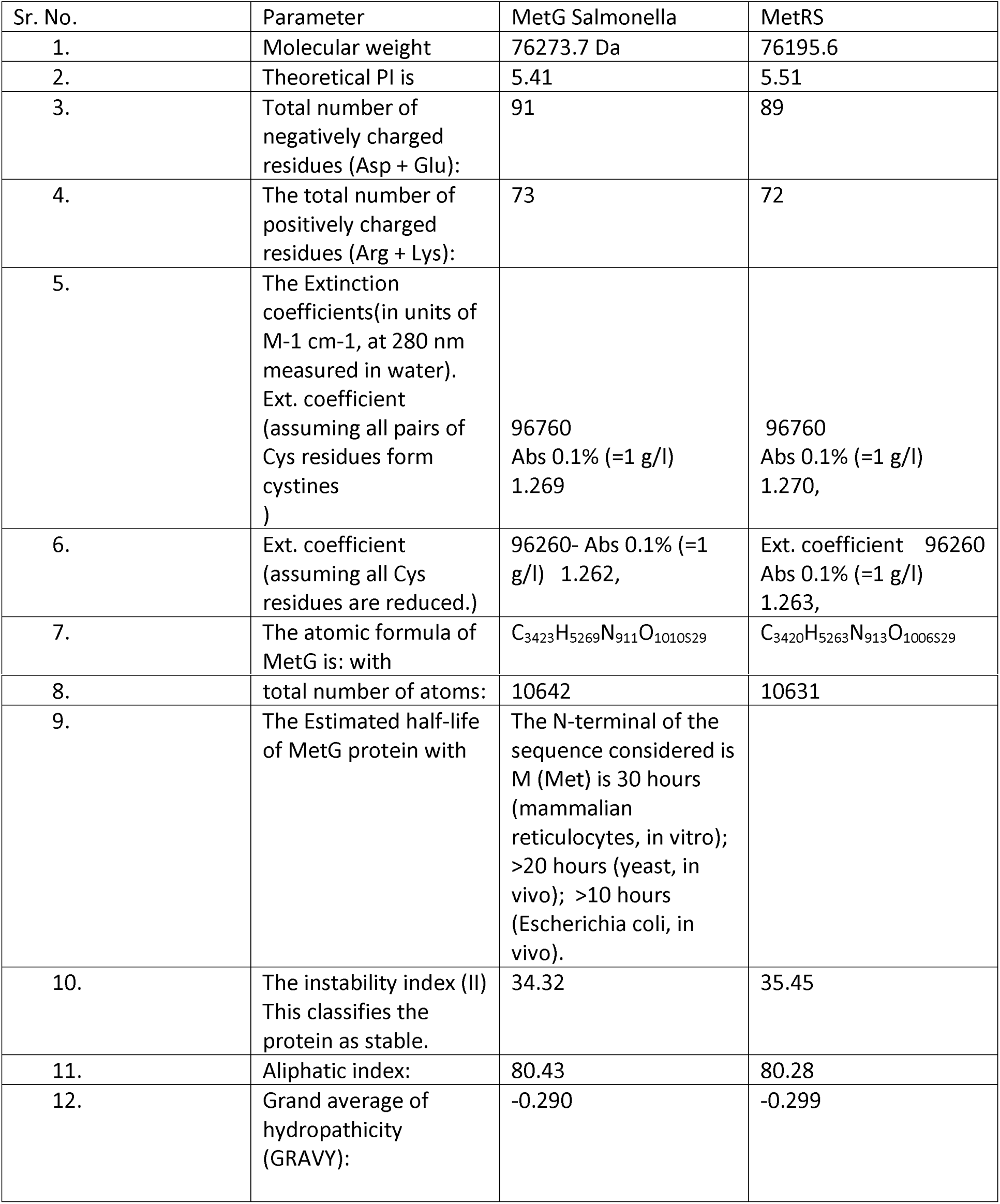
Proto param chart:-

| Sr. No. | Parameter | MetG Salmonella | MetRS |
| --- | --- | --- | --- |
| 1. | Molecular weight | 76273.7 Da | 76195.6 |
| 2. | Theoretical PI is | 5.41 | 5.51 |
| 3. | Total number of negatively charged residues (Asp + Glu): | 91 | 89 |
| 4. | The total number of positively charged residues (Arg + Lys): | 73 | 72 |
| 5. | The Extinction coefficients(in units of M-1 cm-1, at 280 nm measured in water).<br>Ext. coefficient (assuming all pairs of Cys residues form cystines ) | 96760<br>Abs 0.1% (=1 g/l)<br>1.269 | 96760<br>Abs 0.1% (=1 g/l)<br>1.270, |
| 6. | Ext. coefficient (assuming all Cys residues are reduced.) | 96260- Abs 0.1% (=1 g/l) 1.262, | Ext. coefficient 96260<br>Abs 0.1% (=1 g/l)<br>1.263, |
| 7. | The atomic formula of MetG is: with | C<sub>3423</sub>H<sub>5269</sub>N<sub>911</sub>O<sub>1010</sub>S<sub>29</sub> | C<sub>3420</sub>H<sub>5263</sub>N<sub>913</sub>O<sub>1006</sub>S<sub>29</sub> |
| 8. | total number of atoms: | 10642 | 10631 |
| 9. | The Estimated half-life of MetG protein with | The N-terminal of the sequence considered is M (Met) is 30 hours (mammalian reticulocytes, in vitro); >20 hours (yeast, in vivo); >10 hours (Escherichia coli, in vivo). |  |
| 10. | The instability index (II)<br>This classifies the protein as stable. | 34.32 | 35.45 |
| 11. | Aliphatic index: | 80.43 | 80.28 |
| 12. | Grand average of hydropathicity (GRAVY): | -0.290 | -0.299 |

**Table5.** The features of methionyl-tRNAsynthetase are shown in following table.

| Sr. No. | Features | Location/Qualifiers |
| --- | --- | --- |
| 1. | methionyl-tRNA synthetase | 6..677 |
|  | catalytic core domain of methionyl-tRNA | 7..372 |
| 2. | active site | 13..14,16,53,254,257..258,261,298,302 |
| 3. | HIGH motif | 22..25 |
| 4. | KMSKS motif | 333..337 |
| 5. | Anticodon_la_Met | 375..507 |
| 6. | tRNA binding surface [nucleotide binding | 376,383..384,387..388,391..392,395..396,399..400, 453..454,456..457,462 |
| 7. | anticodon binding site | 383,387..388,391..392,395..396,399..400,456,462 |
| 8. | tRNA_bind_EcMetRS_like | 571..677 |
| 9. | dimer interface [polypeptide binding | 572,619,636..638,641,653..656,663,665,667,670, 674..677 |
| 10. | putative tRNA-binding site [nucleotide binding] | 597,609,636,640,647,650 |

Ramachandran Plot (Table 6.) Calculations for 3D model of MetG protein of Salmonella typhimurium indicate Residues in most favoured regions [A,B,L] is higher in MetG than MetRS indicating better model estimation by for MetG compared to MetRS.

**Table 6.** Ramachandran Plot Calculations for 3D model of MetG protein of Salmonella typhimurium

| Ramachandran Plot parameters | MetG Salmonella | MetG Salmonella% | MetGE.coli | MetGE.coli % | QQT | QQT % |
| --- | --- | --- | --- | --- | --- | --- |
| Residues in most favoured regions [A,B,L] | 549 | 91.5% | 534 | 89.6% | 446 | 91.4% |
| Residues in additional allowed regions [a,b,l,p] | 44 | 7.3% | 45 | 7.6% | 41 | 8.4% |
| Residues in generously allowed regions [~a,~b,~l,~p] | 3 | 0.5% | 11 | 1.8% | 1 | 0.2% |
| Residues in disallowed regions | 4 | 0.7% | 6 | 1.0% | 0 | 0.0% |
| Number of non-glycine and non-proline residues | 600 | 100.0% | 596 | 100.0% | 488 | 100.0% |
| Number of end-residues (excl. Gly and Pro) | 2 |  | 2 |  | 2 |  |
| Number of glycine residues (shown as triangles) | 45 |  | 46 |  | 33 |  |
| Number of proline residues | 30 |  | 33 |  | 23 |  |
| Total number of residues | 677 |  | 677 |  | 546 |  |

Based on an analysis of 118 structures of resolution of at least 2.0 Angstroms and R-factor no greater than 20%, a good quality model would be expected to have over 90% in the most favoured regions.

Secondary structure difference- Sequence differences in protein gives rise to minor differences in secondary structures which could be visually observed in Fig.4. No coils, signal peptide and transmembrane protein region was detected for current structures. The visual plus amino acid positional differences which could be visualised by zooming in of image would help to address issue of determining secondary structural differences at that position.

**Fig.4.**
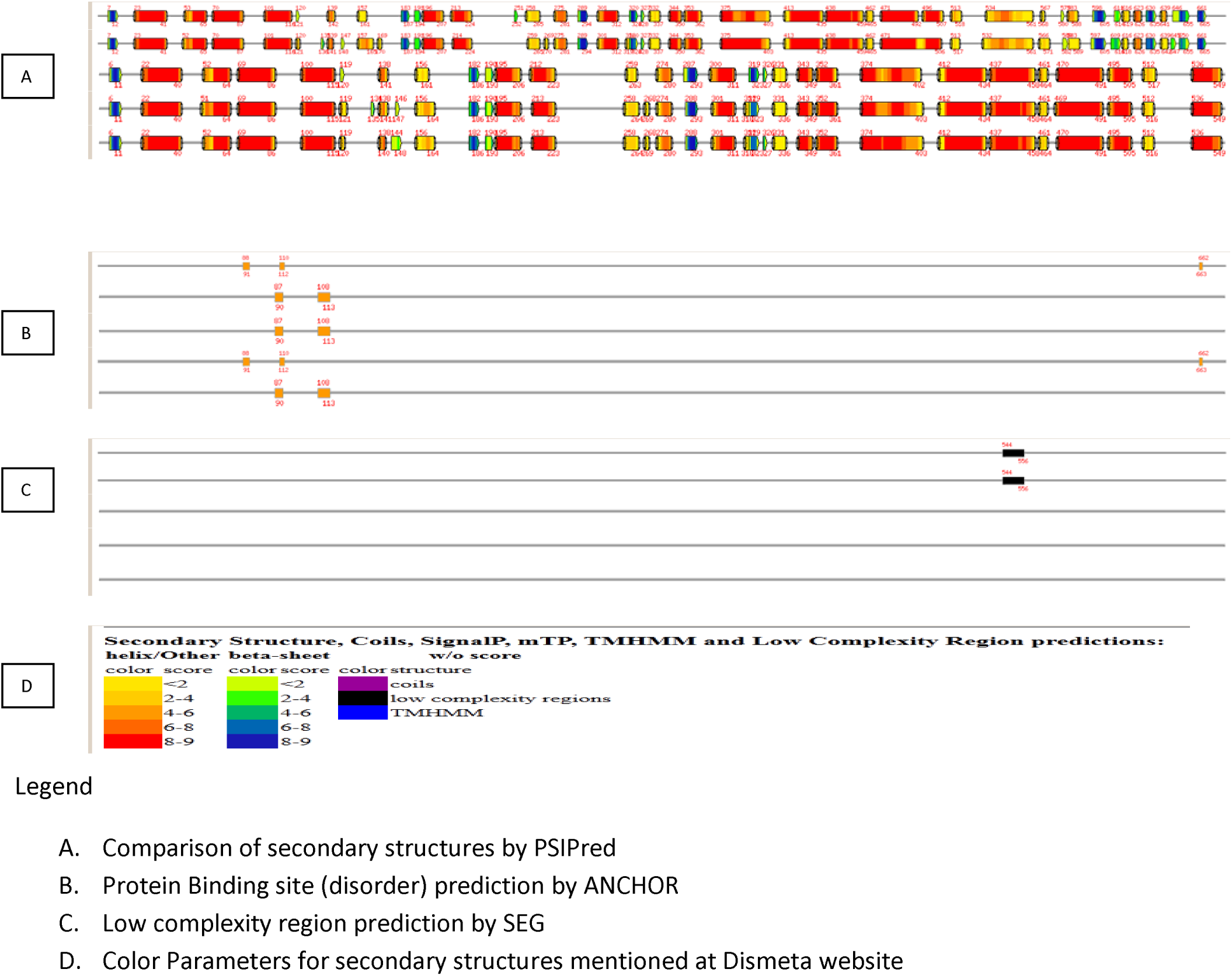
Secondary Structure difference by PSIPred_helix between MetG, MetRS, MetG551, MetRS551 and 1QQT_A respectively in each line

The phylogenetic tree shows close similarity in metRS and MetG551. The metG and metRS551 fall on same root. metG and 1QQT_A found to be more closely related(Fig.6).

### Protein Disorder comparison

The disorder regions predicted by eight methods in Dismeta were compared for MetG, MetRS, MetG551, MetRS551 and 1QQT_A. The graphical presentation helps to understand both positional difference in same length sequences as MetG and MetRS (667bp) or between MetG551, MetRS551 and 1QQT_A (2-552bp). The five structures can be compared by noting positional disorder region differences. Protein disorder difference-Sequences compared for disorder shows differences in individual as well as consensus disorder region. Disorder is more in between 540- 565 and 660-677 Amino acid region. Multiple disorder methods could be compared among themselves visually for same region of given proteins, showing inconsistency in prediction of protein disorder by different methods owing to use of different algorithms by each method (Fig.5).

**Fig.5.**
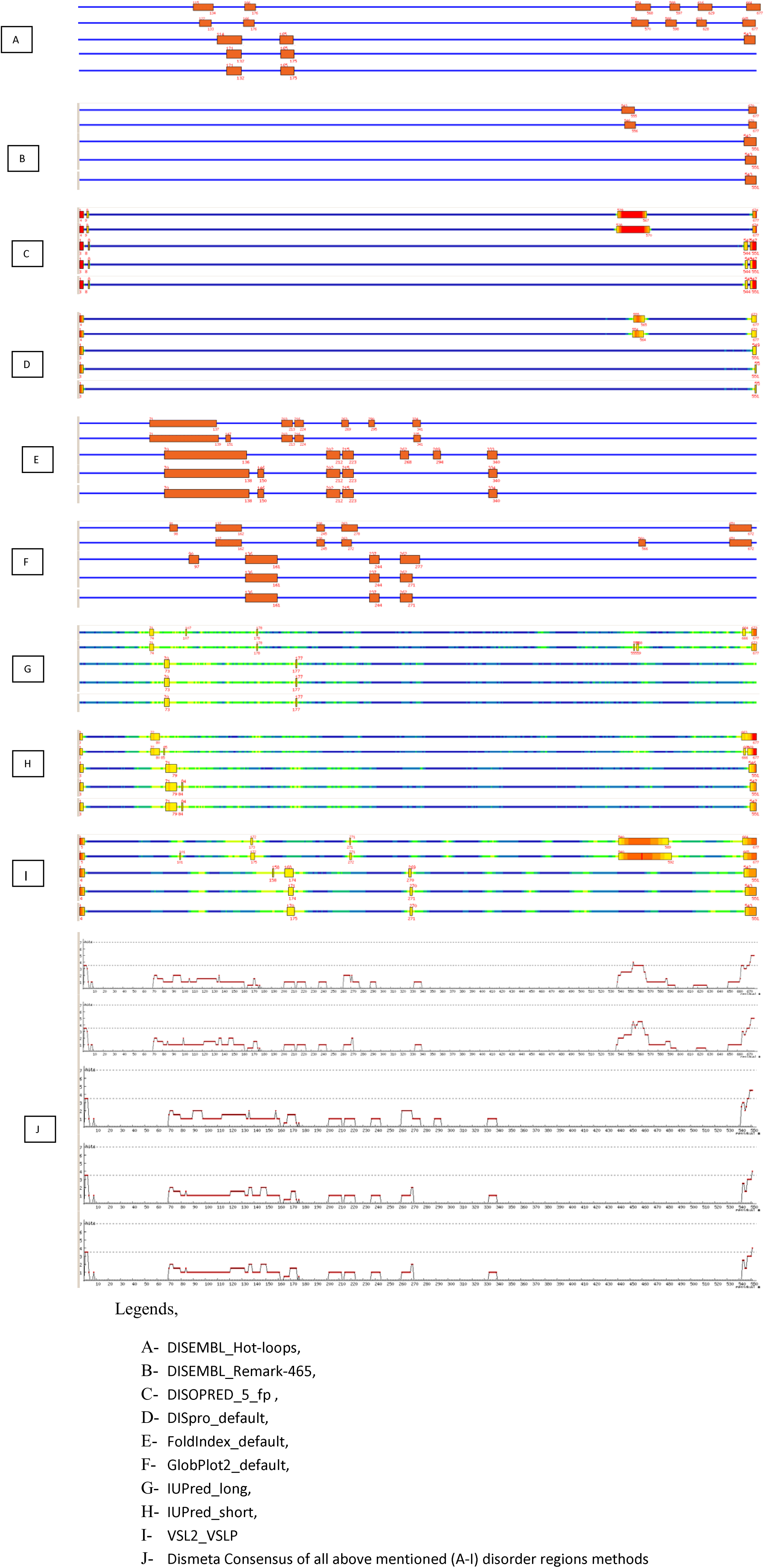
Protein disorder differences between MetG, MetRS, MetG551, MetRS551 and 1QQT_A respectively in each line.

### 3D Structure Validation

Modelled structures were subjected to model validation Table.7, Table. 7b shows higher score of ERRAT for experimental structure than modeled structure. The model validation shows higher score for MetG than MetRS by ERRAT, EVAL123D and VERIFY. Only 1d-3d evaluation score by EVDTREE shows higher score for MetRS compared to metG. ProSA score better (more negativeZ score) for QQT than modeled structures as it is experimental structure (Tabel. 8b). The good score for metG than metRS. So on average model validation quality for MetG is higher than MetRS. Docking shows more tightly binding sites in first three hit (more negative delta G) for metG>metRS> metG551 > metRS551 > QQT. The modeled protein gives different binding site than experimental structure in initial hit in docking, further ability of full length protein for binding with ligand is higher than partial protein. It helps to understand slight structural differences by lower degree of amino acid changes at primary structure level could affect multiple parameters as secondary structures, 3D structure and docking.

**Table7a.**
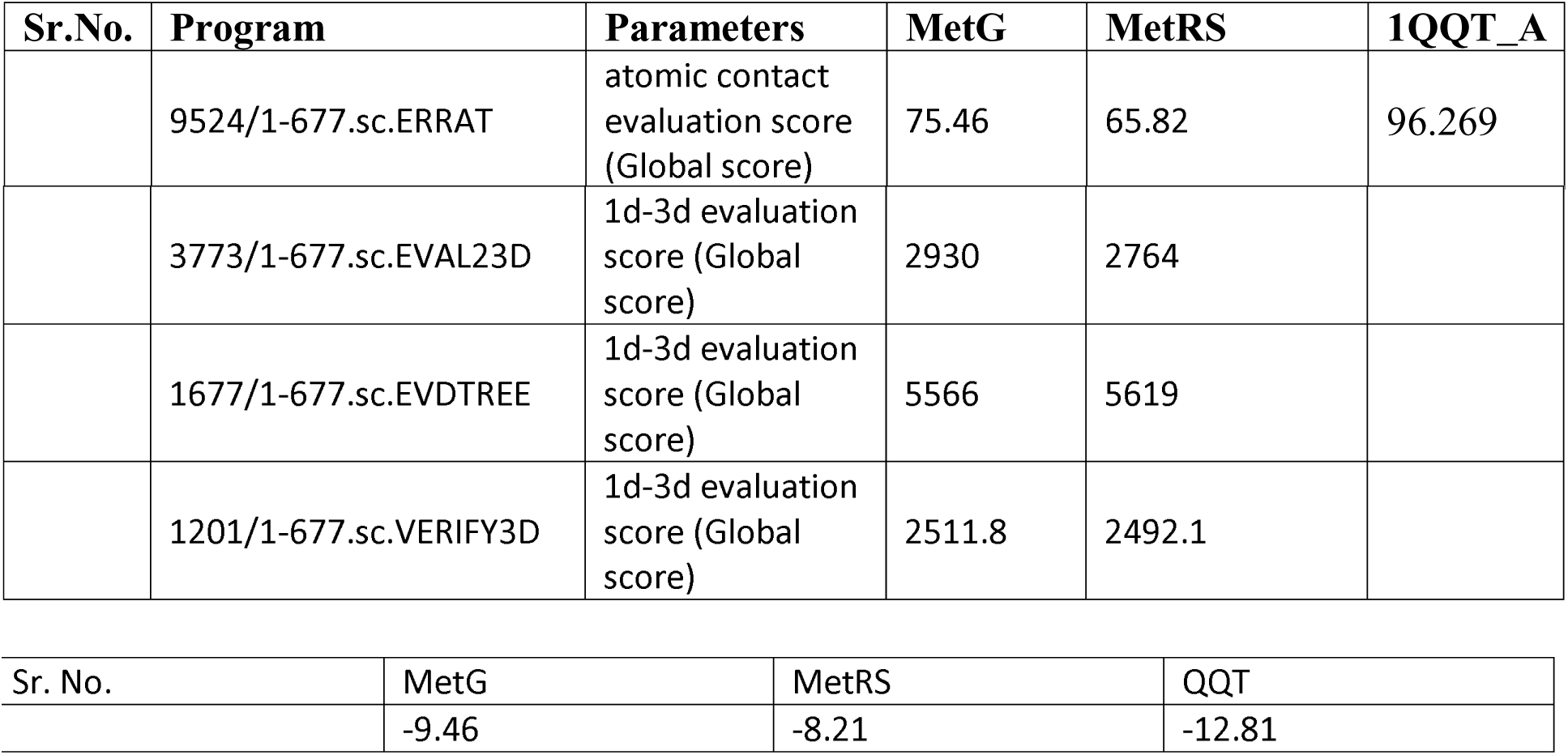
Model Validation-by Protein Analysis ToolKit (PAT) b. ProSA:- Z Score

**Table8a.** SuperPose RMSD analysis chart b. Pairwise structural alignment based on combinatorial extension (CE):-

| Sr. No. | Name of structures Superimposed | Identity | Similarity | Score | Parameter | Local RMSD |  |  |  | Global RMSD |  |  |  |
| --- | --- | --- | --- | --- | --- | --- | --- | --- | --- | --- | --- | --- | --- |
|  |  |  |  |  |  | Alpha Carbons | Back Bone | Heavy | All | Alpha Carbons | Back Bone | Heavy | All |
| 1 | MetG and MetRS | 644/677 (95.1%) | 662/677 (97.8%) | 3419.0 | RMSD | 0.48 | 0.50 | 1.07 | 1.07 | 14.11 | 14.10 | 14.01 | 14.01 |
|  |  |  |  |  | Atoms | 563 | 2252 | 4465 | 4465 | 676 | 2704 | 5307 | 5307 |
| 2 | MetG and 1QQT_A | 520/677 (76.8%) | 534/677 (78.9%) | 2788.0 | RMSD | 0.66 | 0.68 | 1.15 | 1.15 | 0.66 | 0.68 | 1.15 | 1.15 |
|  |  |  |  |  | Atoms | 546 | 2184 | 4341 | 4341 | 546 | 2184 | 4341 | 4341 |
| 3 | MetG, MetRS and 1QQT_A | - | -- | - | RMSD | 0.417 | 0.433 | 0.783 | 0.783 | 0.417 | 0.433 | 0.783 | 0.783 |
| 4 | MetG and MetG | 7/677 (100.0%) | 677/677 (100.0%) | 3560.0 | RMSD | 0 | 0 | 0 | 0 | 0 | 0 | 0 | 0 |
|  |  |  |  |  | Atoms | 677 | 2708 | 5373 | 5373 | 677 | 2708 | 5373 | 5373 |
| 5 | MetG551 and MRS551 | 520/547 (95.1%) | 534/547 (97.6%) | 2791.0 | RMSD | 0.23 | 0.25 | 0.73 | 0.73 | 0.23 | 0.25 | 0.73 | 0.73 |
|  |  |  |  |  | Atoms | 544 | 2176 | 4335 | 4335 | 544 | 2176 | 4335 | 4335 |
| 6 | MetG551 and 1QQT_A | 519/548 (94.7%) | 533/548 (97.3%) | 2784.0 | RMSD | 0.62 | 0.62 | 1.07 | 1.07 | 0.62 | 0.62 | 1.07 | 1.07 |
|  |  |  |  |  | Atoms | 545 | 2180 | 4336 | 4336 | 545 | 2180 | 4336 | 4336 |
| 7 | MRS551 and 1QQT_A | 542/546 (99.3%) | 542/546 (99.3%) |  | RMSD | 0.63 | 0.64 | 1.04 | 1.04 | 0.63 | 0.64 | 1.04 | 1.04 |
|  |  |  |  |  | Atoms | 544 | 2176 | 4367 | 4367 | 544 | 2176 | 4367 | 4367 |
| 8 | MetG551, MRS551 and 1QQT_A | - | - | - | RMSD | 0.287 | 0.297 | 0.59 | 0.59 | 0.287 | 0.297 | 0.59 | 0.59 |

**Fig. 8b.Pairwise structural alignment based on combinatorial extension (CE):-**
| Sr. No | MetG and MetRS | MetG with QQT | MetRS with QQT |
| --- | --- | --- | --- |
| RMSD | 0.52A | 0.66A | 0.59A |
| Z-Score | 8.3 | 8.4 | 8.4 |
| Sequence identities | 93.1% | 95.2% | 99.6% |

Model Validation-by Protein AnalysisToolKit (PAT) shows that ERRAT, EVAL23D and VERIFY3D scores are higher for MetG than MetRS whereas ERRAT is higher for 1QQT_A(Table 8). As Template proteins selected for model building include Sequences from Pyrococcus, Thermococus and E.coli are used in current modelling.

The MetG model shows more Validity score than E.coliMetRS itself though less than experimentally determined 1QQT_A.Table 7. Matching Fold for current MetG model from PROFUNC indicate matching of MetG structure with E.coli Methionine tRNAsynthetase structure.

### Three dimensional structure difference-

The three dimensional structure is compared by ProFunc-Ramachandran plot shows higher residues in most favored region for Salmonella typhimurium than E.coli as 91.5% vs, 89.6. This shows that even the if protein is modeled from available experimental structures; it could give better prediction for model by Procheck than species E.coli which is included in that experiment. MetG on PROFUNC analysis identified fold matching with Salmonella typhimurium MetG Protein sequence all identifying methionyl tRNA synthetase (Table. 2b), the RMSD score is in randge 0.52-0.58 (Laskowski, et al., 2005). The first hit in PROFUNC is 1qqt. So 1qqt is used as reference experimental structure in protein 3D comparison fig.7, Table 8. The modeled structures (MetG and Met) or (metG551 and metRS551) shows lower RMSD than modeled structure with 1QQT_A. modeled structures superimposition with experimental structure shows lower RMSD than individual structures superposing with experimental structure, this is due to above mentioned better RMSD or modeled structures than modeled with experimental decreasing average. Superimposition by Combinatorial extension also shows matching results as superimposition as shown in Table 8.b.

**Fig. 6.**
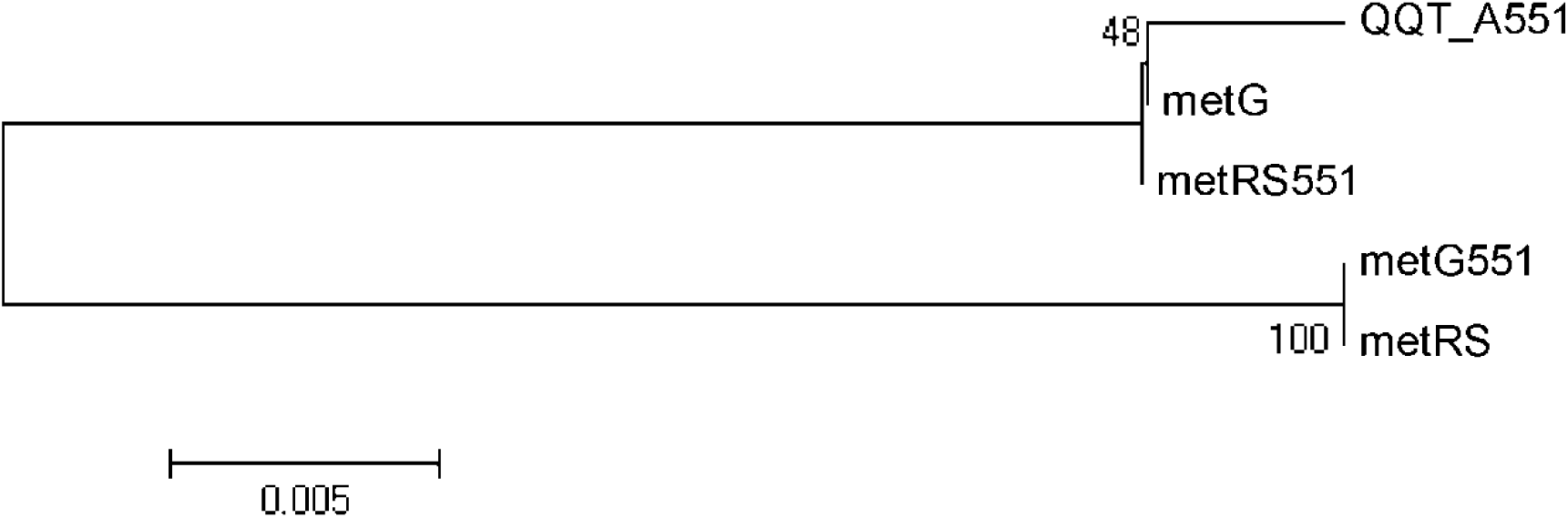
Phylogenetic Analysis by Maximum Likelihood with 500 bootstrap replicate

**Fig7.**
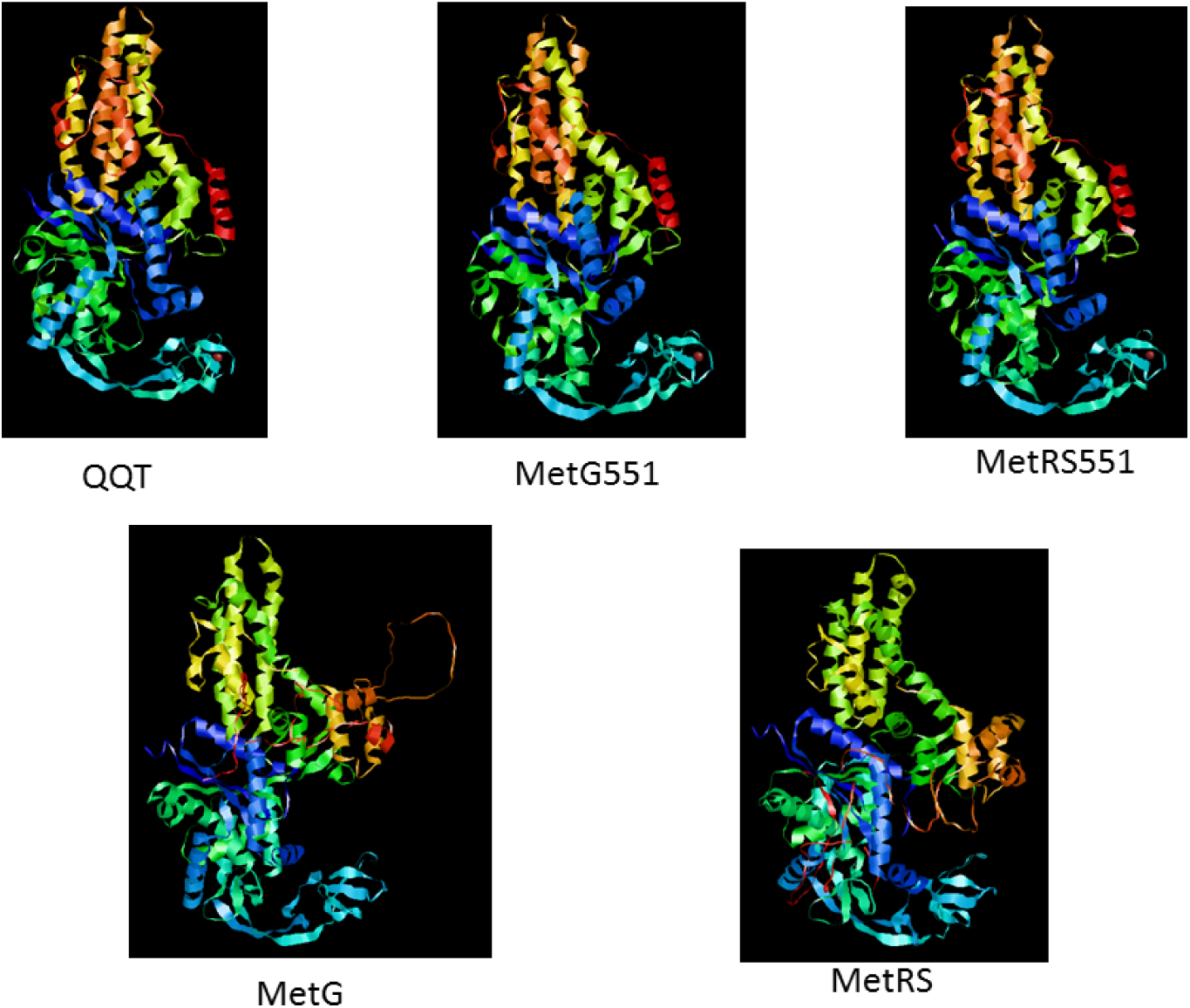
Comparison of Three dimensional structure of MetG, MetRS, MetG551, MetRS551 and 1QQT_A

### Model Comparison

SuperPose RMSD analysis chartTable8a. Indicate MetG and MetRS superimpose has higher local RMSD than Global RMSD. The local and global RMSD are same in MetG and 1QQT_A (Fig.8); MetG, MetRS and 1QQT_A; MetG and MetG. The global RMSD higher than local RMSD in MetG and MetRS indicate efficacy of superimposition in determining differences in Modelled structures. The cause as higher as near 14 RMSD values indicate that the value may be due to prediction error of Modelling as even if 95.1% identity in MetG and MetRS compared to MetG and 1QQT_A in which 76.8% identity where though identity is lower the RMSD value is also lower indicating higher probability of experimental model in superimposition accuracy than in case of modelled proteins. The modelled protein was compared with itself indicating exact similarity owing to RMSD value of 0. The MetG, MetRS and 1QQT_A is lower than (MetG and 1QQT_A) and (MetG and MetRS.

**Fig8.**
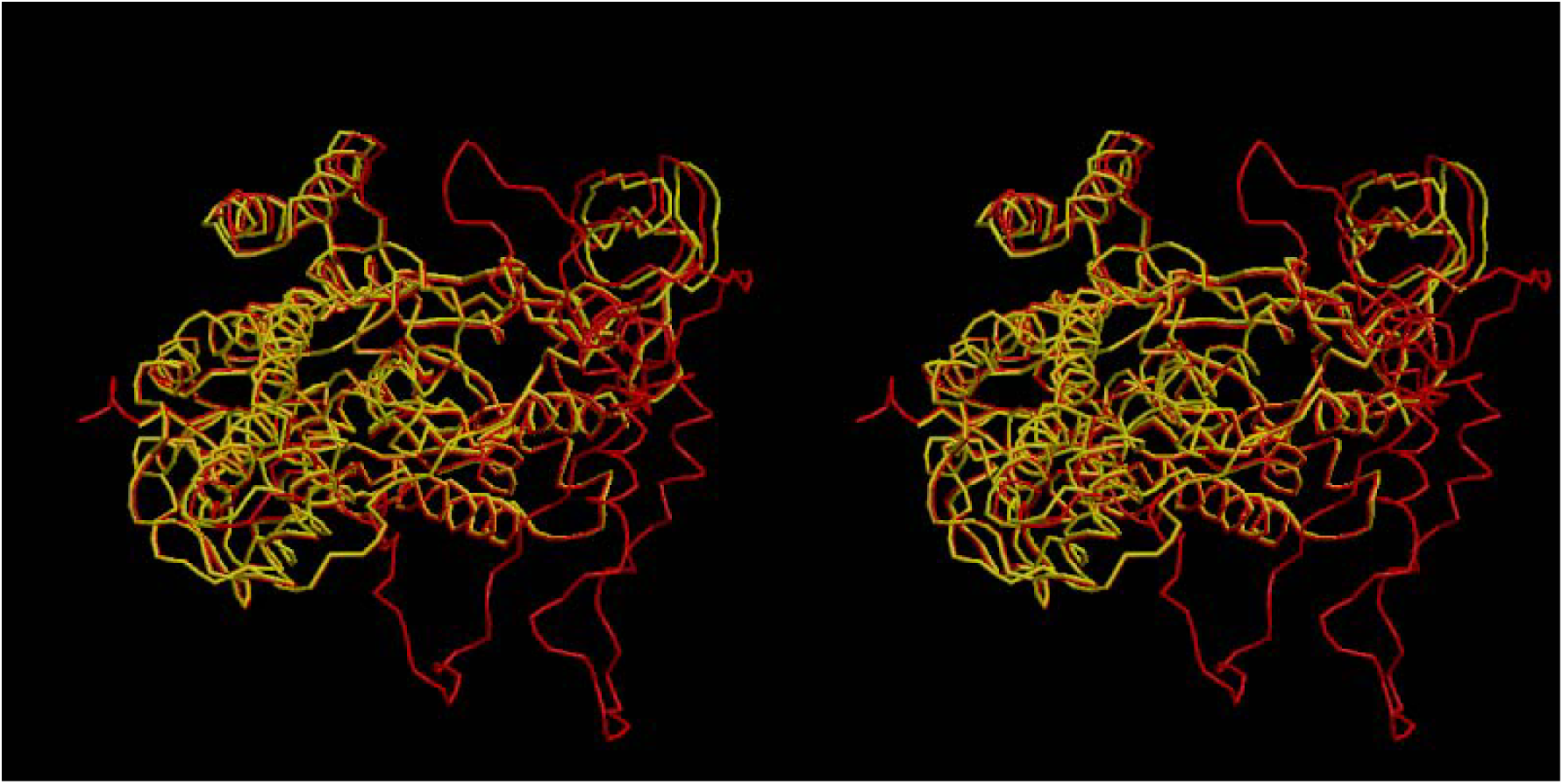
Superimpose of MetG salmonella modeler with QQT

The higher global RMSD in case of MetG-MetRS (both modelled proteins) superimposition indicate necessity of experimental protein for calculating RMSD and reveals variations created during loop modeling (Table8a).

Local RMSD values for all atoms are in following order for superimposed structures MetG and MetG<MetG, MetRS and 1QQT_A <MetG and MetRS<MetG and 1QQT_A(Table8a).

### Swiss Docking

The docking results for each protein with L-Methionine is shown in (table 9.)

**Table 9.**
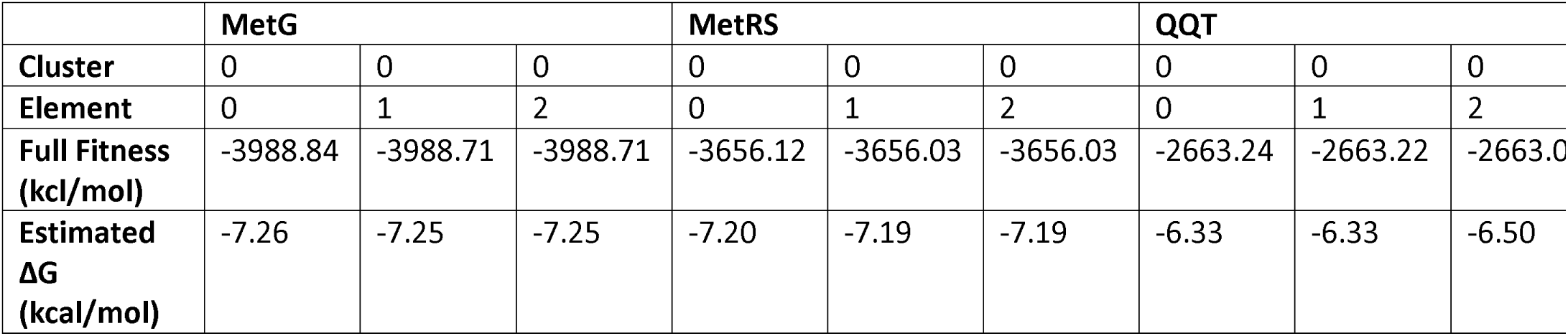
Prediction of Binding Sites. Predicted binding modes for Methionyl-tRNAsynthetase with Methionine

## Discussion

We had compared primary, secondary and tertiary structure of MetG from E.coli and Salmonella spp to find out the impact of subtle differences in the primary sequences on secondary and tertiary structure. The similarity in primary sequence between the two sequence studied was 97.8%. None of the differences in the amino acid sequences was in the functional region of the proteins thus indicating that functional region are important for MetG activity and are conserved. MetG is an important enzyme involved in the charging of tRNA with an amino acid for translation. There is only one copy of MetG in the genome of these bacteria and thus conservation of the activity of this enzyme is important for these bacteria. MetG is also survival gene, knocking it out makes the bacteria unable to survive. Thus it can be assumed that the difference in the amino acid sequences observed between the two sequences is those amino acids which are not important for the function of this protein. When the secondary structure of these two proteins was compared, a slight difference in the secondary structure of the MetG of the two species was observed. It can be further assumed that this difference is not significant for the activity of this enzyme since the known functional region of these proteins is conserved.

Analysis of three dimensional structures shows a RMSD difference of 0.2-0.6 (Table8) whether full length or partial structure are compared. The difference in RMSD below 0.5 indicates close similarity in 3D structure, observed difference in RMSD of 0.2-0.6 indicate that two structures have differences in 3D structure despite high sequence similarity; the reason for that could be due to difference in secondary structure of these proteins (Fi4, Fig.5), which are getting reflected in three dimensional structures.

To check if the protein retains functional activity despite differences in 3D structure, docking with one of the two ligands was done. Docking data indicate insignificant difference in affinity of methionine for the two proteins (MetG and MetRS ΔG -7.26 and -7.20); (QQT, MetG551, MetRSΔG -6.33, -8.19, -7.28) indicating that the protein retains functional activity. But the creation of additional binding sites (Table 9) on full length sequence modeling compared to partial sequence model indicate that the difference can be correlated with the protein disorder region in full length sequence (Fig.5). But as both protein sequences were taken from non mutant protein sequences these differences in 3D structures are not affecting functional activity. The phylogenetic analysis of closely related structure gives less information about relationship as metG and metRS551 fall on same root despite they are from different bacteria (Salmonella Sp and E.Coli) (Fig.6).

A study carried out with homology modeled 3 D structure and docking when subsequently tested with crystal structure and docking revealed almost identical result (Carlsson, et al., 2011). In this study we have modeled 3D structure from crystal structure of a very closely related (97.8% sequence similarity) protein, and therefore assumed that the modeled structure would resemble the crystal structure to a large extent.

Our results indicate that subtle differences in amino acid sequences are reflected in the secondary and tertiary structure of a protein without affecting functional activity. The methodology we had adopted can be used for in silico study of primary sequence difference on functional activity of proteins.

## Conclusion

The nucleotide level non-silent mutation gives rise to amino acid sequence differences and how to know effect of these sequence disparity on secondary structure and 3D structure with functional level differences is burning area of structural bioinformatics helping to elucidate actual cell level phenomenon of such mutations. Current Research flow gives methodological bioinformatics approaches to study amino acid differences at various levels.

## Acknowledgement

We like to thank Indian Veterinary Research Institute, Indian Council of Agricultural Research (ICAR) and Council for Scientific and industrial Research (CSIR) for supporting this work.

